# Molecular insights into *Vibrio cholerae*’s intra-amoebal host-pathogen interactions

**DOI:** 10.1101/235598

**Authors:** Charles Van der Henst, Stéphanie Clerc, Sandrine Stutzmann, Candice Stoudmann, Tiziana Scrignari, Catherine Maclachlan, Graham Knott, Melanie Blokesch

## Abstract

*Vibrio cholerae*, which causes the diarrheal disease cholera, is a species of bacteria commonly found in aquatic habitats. Within such environments, the bacterium must defend itself against predatory protozoan grazers. Amoebae are prominent grazers, with *Acanthamoeba castellanii* being one of the best-studied aquatic amoebae. We previously showed that *V. cholerae* resists digestion by *A. castellanii* and establishes a replication niche within the host’s osmoregulatory organelle. In this study, we deciphered the molecular mechanisms involved in the maintenance of *V. cholerae*’s intra-amoebal replication niche and its ultimate escape from the succumbed host. We demonstrated that minor virulence features important for disease in mammals, such as extracellular enzymes and flagellum-based motility, play a key role role in the replication and transmission of *V. cholerae* in its aqueous environment. This work, therefore, describes new mechanisms that provide the pathogen with a fitness advantage in its primary habitat, which may have contributed to the emergence of these minor virulence factors in the species *V. cholerae.*

---

The severe diarrheal disease cholera is not extinct. Seven cholera pandemics have been recorded in modern history, most notably in developing countries, and the latest is still ongoing^1,2^. An estimated three million new cases of cholera occur every year leading to almost 100,000 annual deaths^3,4^. Cholera is caused by ingestion of the bacterium *Vibrio cholerae.* Toxigenic strains of this species are capable of damaging the host due to the presence of so-called virulence factors, which refers “to the elements (i.e. gene products) that enable a microorganism to colonize a host niche where the organism proliferates and causes tissue damage or systemic inflammation.”^5^ Bacterial strains without these factors are usually attenuated with respect to the infection process (in human or animal models).

For *V. cholerae*, the two major virulence factors, cholera toxin and the toxin-coregulated pilus, play a pivotal role in the infection process, but additional minor virulence factors have also been identified. These include factors such as outer membrane porins, a pore-forming hemolysin, diverse proteases, N-Acetyl-glucosamine binding protein (GbpA), flagellum-based motility, to name a few^6-9^. The hemagglutinin/protease (HA/P or HapA), for example, is a zinc-metalloprotease^10^, which was first identified due to its mucinase activity^11^. This enzyme was later demonstrated to not only cause hemagglutination but to also hydrolyze fibronectin, mucin, and lactoferrin, all of which were thought to contribute to the host defense against *V. cholerae*^12^. HapA also causes cell rounding, loss of the barrier function of the epithelial layer, and, ultimately, detachment of the cells under tissue culture conditions^13,14^. Consistent with these *in vitro* activities, a 10-fold increase in the 50% lethal dose (LD_50_) in the absence compared to the presence of the HapA protease for *V. cholerae* strains that otherwise lack cholera toxin was reported^15^. It was therefore suggested that HapA participates in the attachment as well as in the detachment of *V. cholerae* from the gut epithelium^12,16^ and that tissue destruction through HapA-mediated cleavage of surface-exposed proteins may contribute to fluid leakage and the stimulation of proinflammatory responses^7^.

A second well-characterized minor virulence factor that is widespread among *Vibrio* species^17^ is hemolysin (HlyA), which is also known as *V. cholerae* cytolysin (VCC), β-hemolysin, and vacuolating toxin. Hemolysin is a secreted protein belonging to the family of pore-forming toxins, as it forms aqueous channels *in vitro* in host cells with an estimated diameter of 0.7nm^18^. *In vivo* data by Ichinose and colleagues showed that purified hemolysin induced intestinal fluid accumulation in rabbits and in orally inoculated suckling mice^19^, two commonly used animal models of cholera. The authors therefore concluded that hemolysin was an enterotoxic factor that contributed to gastroenteritis caused by non-pandemic *V. cholerae* strains (e.g., non-O1 serogroup strains)^19^. The role of HlyA as enterotoxic or diarrheagenic factor, especially in vaccine strains lacking cholera toxin, was later confirmed by Alm *et al.* who also demonstrated that mice infected with hemolysin-deficient *V. cholerae* strains survived longer than those inoculated with their hemolysin-positive parental strains^20^. More recent studies using streptomycin-treated adult mice showed that hemolysin was the major cause of lethality in this animal model of cholera^21^. Moreover, it has been suggested that HlyA and other secreted accessory toxins modify the host environment, thereby contributing to prolonged disease-free colonization that could contribute to disease transmission via asymptomatic carriers^22^.

It was proposed that *hlyA* might be located on a pathogenicity island due to the close proximity of additional genes that likewise encode extracellular enzymes^23^. This region encodes apart from hemolysin (*hlyA*) a lipase (*lipA*), a metalloprotease (*prtV*), and a lecithinase/phospholipase (*lec;* also known as thermolabile hemolysin). The lecithinase/phospholipase-mediated activity of the latter enzyme was first characterized by Felsenfeld^24^ and later again by Fiore *et al.*^25^, but molecular details and its concrete function are still unknown.

To date, a limited number of studies have shown that minor virulence factors, such as those mentioned above, are also advantageous in an environmental context^26,27^. Indeed, despite *V. cholerae* being an aquatic bacterium that is well adapted for survival in this environment^28^, the molecular details about *V. cholerae*’s environmental lifestyle are still lacking. To address this knowledge gap, we aimed to elucidate the molecular mechanisms that *V. cholerae* would use as part of its environmental lifestyle, such as its interactions with grazing amoebal predators. In this context, we recently demonstrated that *V. cholerae* survives predation by *Acanthamoeba castellanii*, an aquatic species of amoebae, through the evolution of two distinct phenotypes. Firstly, upon phagocytosis by the feeding amoebal trophozoite, a fraction of the ingested *V. cholerae* population can resist amoebal digestion, then exit the phagosomal pathway by exocytosis. Notably and in contrast to a previous study^29^, we never observed free *V. cholerae* cells in the cytosol of such trophozoites^30^, indicating that the bacteria are not released from food vacuoles intracellularly. Secondly, the pathogen colonizes the amoeba’s contractile vacuole, which is its osmoregulatory organelle and, therefore, of prime importance for the survival of *A. castellanii.* We showed that the bacterium proliferates within this replication niche, even upon amoebal encystement, before it ultimately lyses the host^30^. Here, we address the underlying molecular mechanisms of this host-pathogen interaction. We show that *V. cholerae* fine-tunes extracellular enzymes to avoid premature intoxication of its host, allowing the bacterium to take full advantage of the intra-amoebal replication niche for undisturbed and non-competitive proliferation before finally lysing its amoebal host. We also demonstrate that flagellum-based motility is of prime importance for *V. cholerae* to ultimately escape the ruptured host niche to return to the aquatic environment.

## Results

### Visualization of colonized contractile vacuoles by correlative light and electron microscopy

Using confocal laser scanning microscopy, we recently demonstrated that *V. cholerae* could enter the contractile vacuole of *A. castellanii* and replicate within this niche^30^, though underlying molecular mechanisms of this host-pathogen interaction remained largely unknown. This is partially due to the low throughput of transmission electron microscopy (TEM), which only allowed for the imaging of hyper-colonizing *V. cholerae* mutant strains, due to their lack of quorum-sensing^30^, while spatial information on wild-type (WT) bacteria within the amoebal vacuoles remained missing. To circumvent this problem, we developed a correlative light and electron microscopy (CLEM) imaging strategy, which allowed us to study this specific host-pathogen interaction for WT *V. cholerae* for the first time at high resolution. We identified *V.* cholerae-infected *A. castellanii* cells by pathogen-produced green fluorescent protein GFP and then targeted these cells for TEM. Using this method allowed us get information on the morphology of the colonizing bacteria and their spatial distribution. As shown in Fig. 1, the morphologies of such intra-vacuolar WT bacteria possessed the typical *Vibrio* shape. Moreover, the membrane surrounding the contractile vacuole was largely intact, which was consistent with the non-encysted phenotype of the amoeba (Fig. 1a).

**Figure 1.**
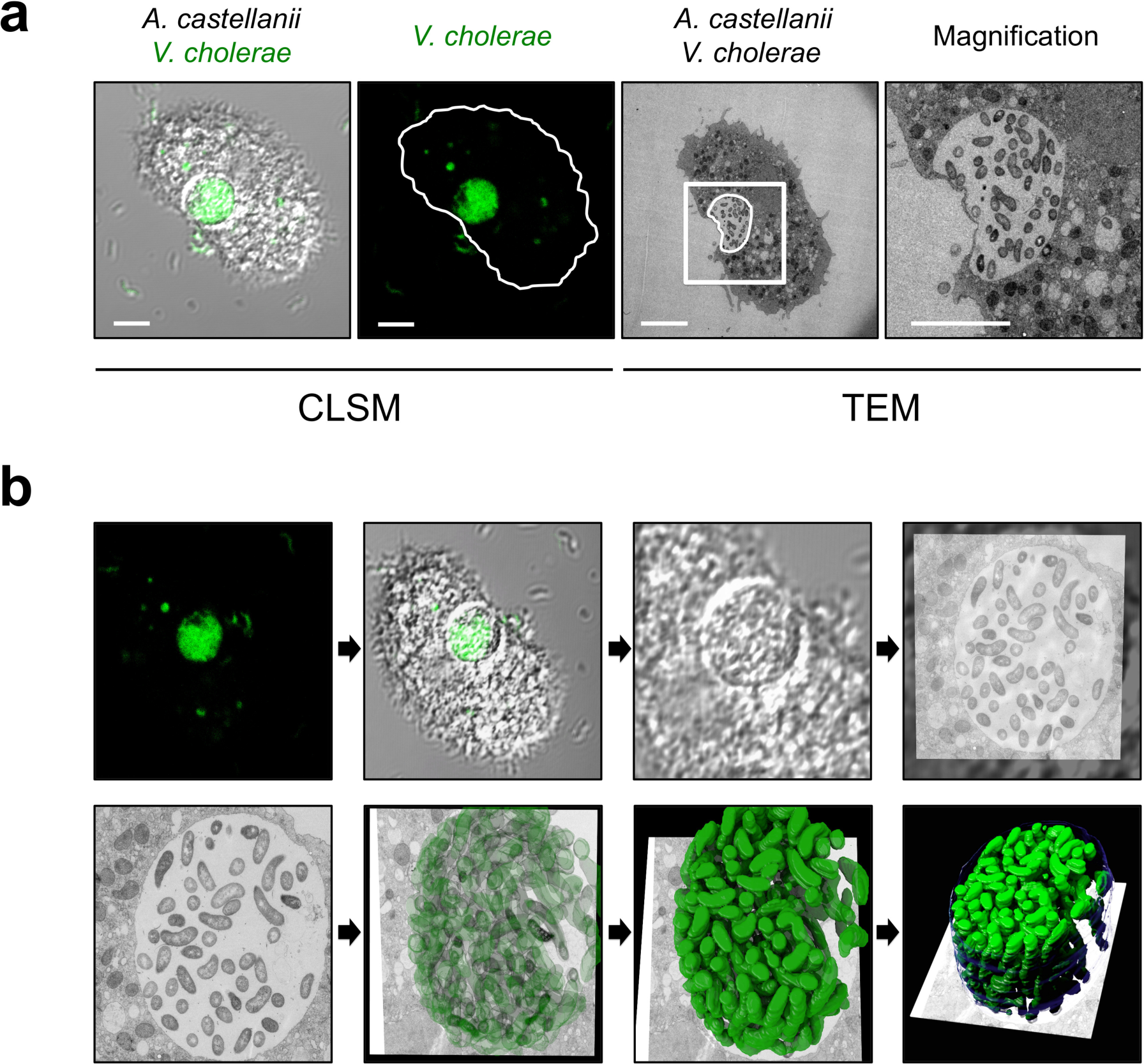
Correlative light and electron microscopy (CLEM) for visualizing wild-type *V. cholerae* inside the contractile vacuole. **(a)** Low- and high-resolution imaging of an infected amoeba. GFP-tagged *V. cholerae* were seen to be localized inside of a contractile vacuole of *A. castellanii* using confocal laser scanning microscopy (CLSM; low resolution) in fixed samples. Shown are a merged image of the transmitted light channel and the green channel (left) and the green channel image alone (second from left). After staining of the sample, the same amoeba was imaged at high resolution using transmission electron microscopy (TEM; right images). Scale bar in all images: 5 μm. **(b)** 3D reconstruction of the colonized contractile vacuole. The region containing the amoeba shown in panel (a) was serially thin sectioned (50 nm thickness) and serial images were taken with the TEM. These images were then aligned to generate a 3D model of the colonized amoeba. Shown are snapshots of the resulting 3D reconstruction movie **(Movie S1)**.

We estimated the number of objects – in this case, bacteria – in the contractile vacuole (Fig. 1b and Movie S1) using an unbiased stereological approach^31^. To do this, a representative WT *V. cholerae*-colonized amoeba, at 20 hours post-primary contact (p.p.c.), was identified in the fluorescent channel after fixation, and, after staining and resin embedding, it was then serially thin sectioned into 50 nm-thick slices (55 slices in total). We then took several dissector pairs, each pair consisting of a reference and a lookup section, and used these to estimate the numbers of bacteria in each. This gave a mean value of 1.35 (± standard deviation of 0.42) for each dissector pair. Given the approximate total volume of the contractile vacuole of 143μm^3^, based on the assumption of a spherical shape, the total number of *V. cholerae* cells was estimated to be 194. This number, and the apparent density of bacteria in the 3D reconstruction (Fig. 1b and Movie 1), correlates well with the active growth of *V. cholerae* in this organelle, as the vacuole’s initial colonization only involved one or a few bacteria^30^. While this experimental approach was elaborate and, therefore, not applicable to large numbers of individual amoeba, an average number of 150-200 *V. cholerae* cells per organelle is also theoretically supported given an average volume of ~μm^3^ (www.BioNumber.org) and 150μm^3^ (based on a maximum diameter of up to 6.6 μm^32^) for bacteria and the contractile vacuoles of *A. castellanii*, respectively.

### The absence of *V. cholerae*’s hemagglutinin protease leads to aberrant amoebal morphologies

As the 3D reconstruction (Fig. 1b and Movie S1) suggested that *V. cholerae* cells were densely packed within the vacuole at this later point of the infection, we wondered how the bacteria eventually escaped the vacuole after amoebal encystation and how the timing was properly regulated. To answer these questions, we first took an educated guess approach and tested a few diverse knockout mutants of *V. cholerae* for their intra-amoebal behavior. We were especially interested in strains that lacked certain extracellular enzymes, as some of those were known to be important in the pathogen’s intra-human lifestyle and its transmission to new hosts.

We therefore challenged *A. castellanii* with a HapA-deficient *V. cholerae* strain (Table S1) and compared the amoebal response to the WT-challenged condition. By doing this, we observed that a vast majority (79.8%) of infected amoebal cells showed an aberrant morphology upon co-incubation with the *hapA*-minus strain for 20 hours, which occurred significantly less often for WT-infected amoebae (11.6%, Fig. 2a and b). These morphological abnormalities ranged from a shrinking or compacting phenotype towards pseudopodia retraction and detachment, all of which ultimately abolished amoebal grazing (Fig. 2 and Fig. S1). Complementation assays, involving a genetically engineered derivative of the *hapA*-minus strain with a new copy of *hapA* on the large chromosome (Fig. S2), were performed using the amoebal infection protocol and fully restored the WT-infected amoebal morphologies (Fig. 2a and b).

**Figure 2.**
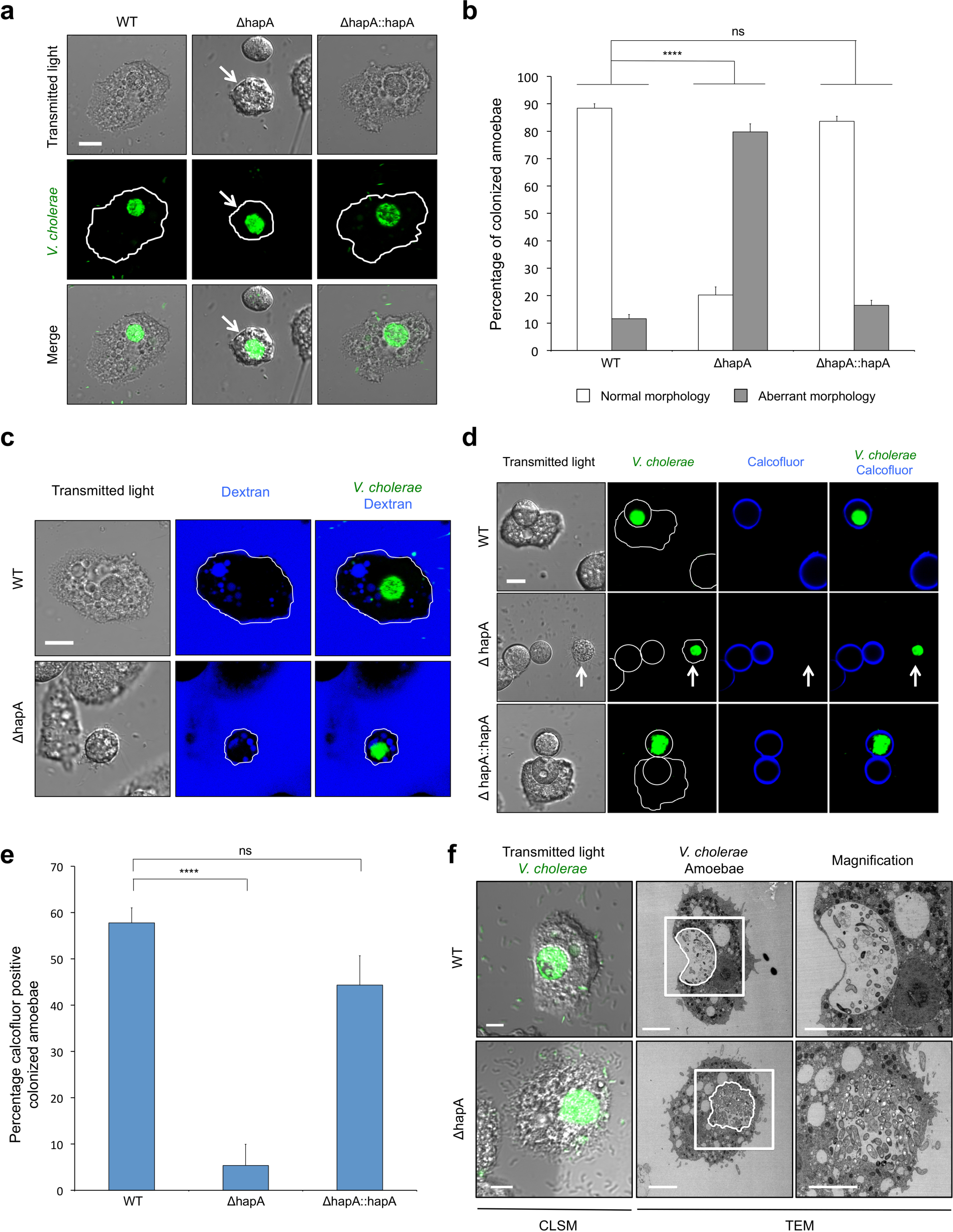
Hemagglutinin protease-deficient *V. cholerae* strains intoxicate amoebal trophozoites. **(a)** CLSM images of amoebae that were colonized by GFP-tagged wild-type (WT), *hapA*-minus (△hapA), or the *hapA* complemented (△hapA::hapA) *V. cholerae* strains at 20 hours p.p.c. Shown are the transmitted light channel (top), the green channel (middle), and the merged image of both channels (bottom). The white arrows highlight the aberrant morphology of the intoxicated amoeba. **(b)** Quantification of normal and aberrant morphologies of amoeba colonized with any of the three bacterial strains mentioned in panel (a). **(c)** Protease-deficient *V. cholerae* mutants do not co-localize with dextran-labeled digestive vacuoles. Labeling of the endosomal pathway using fluorescent dextran shows the residence of WT (top) and ΔhapA (bottom) strains inside the dextran-negative contractile vacuole. **(d)** Cellulose is not deposited around amoebae infected with *hapA*-minus bacteria at 30 hours p.p.c. Amoebae infected with WT, ΔhapA, or the *hapA* complemented strain (ΔhapA::hapA) were stained with calcofluor to visualize deposited cellulose around the cysts. The white arrows point towards a cellulose-deficient *A. castellanii* infected by the hapA- minus strain. **(e)** Quantification of the calcofluor-stained colonized amoebae. Bacterial strains correspond to those in panel (d). **(f)** CLEM images of WT- and *hapA*-minus-infected amoebal trophozoites (details as in Fig. 1). Scale bars: 10 μm (panels a, c, d) and 5 μm (panel f). All graphs show the averages of three independent biological replicates (± SD, as shown by the error bars). Statistics are based on a one-way ANOVA. *** *p* ≤ 0.001; ns (not significant), *p* > 0.05.

Seeing these striking differences between WT-infected and *hapA*-minus-infected amoebae, we wondered whether the latter might cause “amoebal constipation”, meaning the accumulation of undigested phagosomal *V. cholerae* without efficient exocytosis. Indeed, based on the confocal microscopy images, we could localize the accumulated bacteria due to their green fluorescence. However, recognition of the contractile vacuole in the transmitted light channel was often difficult for *hapA*-minus mutant-infected amoebae due to their compaction and malformation (Fig. 2a). Therefore, it seemed possible that the bacteria were contained in digestive food vacuoles of the endosomal pathway, blocking amoebal digestion as opposed to being localized to the contractile vacuole, as is the case for WT *V. cholerae.* To distinguish between both scenarios (localization within a digestive vacuole or within the contractile vacuole), we labeled the amoebal endosomal pathway with fluorescently labeled dextran. As the dextran- and bacteria-derived fluorescent signals did not overlap, these experiments confirmed the intra-contractile vacuolar localization of the *hapA*-minus strain and excluded its massive accumulation within dextran-labeled digestive food vacuoles in intoxicated amoebae (Fig. 2c).

### HapA-deficient *V. cholerae* impair amoebal encystation and cause vacuolar abnormalities

Next, we wondered whether, despite these unusual amoebal morphologies, the life cycle between *hapA*-minus *V. cholerae* and *A. castellanii* would proceed as previously reported for the WT^30^. Such a progression included amoebal encystation followed by the escape of *V. cholerae* from the contractile vacuole into the cyst’s cytosol, leading, ultimately, to lysis of the amoebal host. To determine this, we imaged infected amoebae at a later time point (30 hours p.p.c) to allow the life cycle progression to develop. To better distinguish aberrantly rounded cells from cysts, we used calcofluor staining, as this fluorescence stain is known to bind to the deposited cellulose layer in *Acanthamoeba* cysts^33^. As shown in Fig. 2d (and quantified in Fig. 2e), the ability to form cellulose-positive cysts was severely impaired in amoebae infected with the *hapA*-minus mutant strain. This impaired encystation contrasted greatly to that of both the WT-infected *A. castellanii* and the complemented *hapA*-minus strain (Fig. 2d and e). We therefore concluded that the absence of the HapA protease leads to premature amoebal intoxication and, consequently, a defect in the encystation process.

To better understand the aberrant morphologies and the defect in the encystation process of the amoebae infected by the mutant strain, we again used CLEM to obtain high-resolution images of colonized amoebae. Impaired membrane integrity around the contractile vacuole in amoeba infected with *hapA*-minus *V. cholerae* was seen from the TEM images (Fig. 2f). The WT-infected contractile vacuoles seemed to maintain their internal pressure, leading to a clear separation between the content of the contractile vacuole and the amoebal cytosol (Fig. 1a and 2f), though this evident distinction was lacking in host cells that were infected by the mutant *V. cholerae* strain (Fig. 2f).

### Uncontrolled hemolysis leads to premature amoebal intoxication

We speculated that the seemingly collapsed contractile vacuole membranes might have been due to amoebal membrane rupture caused by the *hapA*-minus mutant strain. Given that bacterial pathogens often cause membrane rupture through the secretion of pore-forming toxins^34^, we wondered whether the secreted hemolysin of *V. cholerae* (HlyA) might be involved in the observed amoebal intoxication. Consistent with this idea was a study by Tsou and Zhu that showed that the HapA protease degrades HlyA in an *in vitro* assay^35^. This led us to hypothesize that HapA protease-deficient strains of *V. cholerae* would display enhanced hemolysis, which, ultimately, could cause the observed amoebal intoxication. To test this hypothesis, we generated several *V. cholerae* strains that lacked *hapA, hlyA*, or both genes simultaneously (Table S1) and tested them for hemolysis and proteolysis on blood and milk agar plates, respectively. As shown in Fig. S2, we observed a significant correlation between the absence of HapA-mediated protease activity and the presence of hemolysis. There was a strong increase in hemolysis in strains lacking *hapA*, while the absence of *hlyA* fully abolished this activity. Complementation assays, in which the mutant strain contained a new copy of the missing gene elsewhere on the chromosome (Table S1), restored the system to that of the WT (Fig. S2a and b). Interestingly and consistent with the extracellular localization of both enzymes, we showed that the secreted HapA protease from a WT strain was also able to inactivate HlyA that was released by a co-cultured *hapA*-minus mutant strain (Fig. S2). However, when we engineered an *hlyA*-overexpression strain (Table S1 and Fig. S2), the protease activity exerted by the strain itself or from the co-cultured WT bacteria was insufficient to abolish hemolysis (Fig. S2).

With these newly constructed strains in hand, we then tested their effect on *A. castellanii.* Consistent with our hypothesis that a *hapA* mutant of *V. cholerae* would possess enhanced hemolysin activity, which, ultimately, would result in amoebal intoxication, we found that the absence of *hlyA* in the protease-minus background restored normal host morphology and allowed cyst formation comparable to that observed for WT infection (Fig. 3a and b and Fig. S3a to d). Infection of *A. castellanii* by the *V. cholerae* double mutant (ΔhapAΔhlyA) also resulted in an intact integrity of the contractile vacuole membrane (Fig. 3c and Fig. S3e). Overproduction of HlyA in a WT background strain, however, fully phenocopied or even aggravated the protease-minus phenotypes (Fig. 3a and b and Fig. S3a to d). These data, therefore, support the notion that HlyA is causing amoebal intoxication and that the HapA protease counteracts this effect.

**Figure 3.**
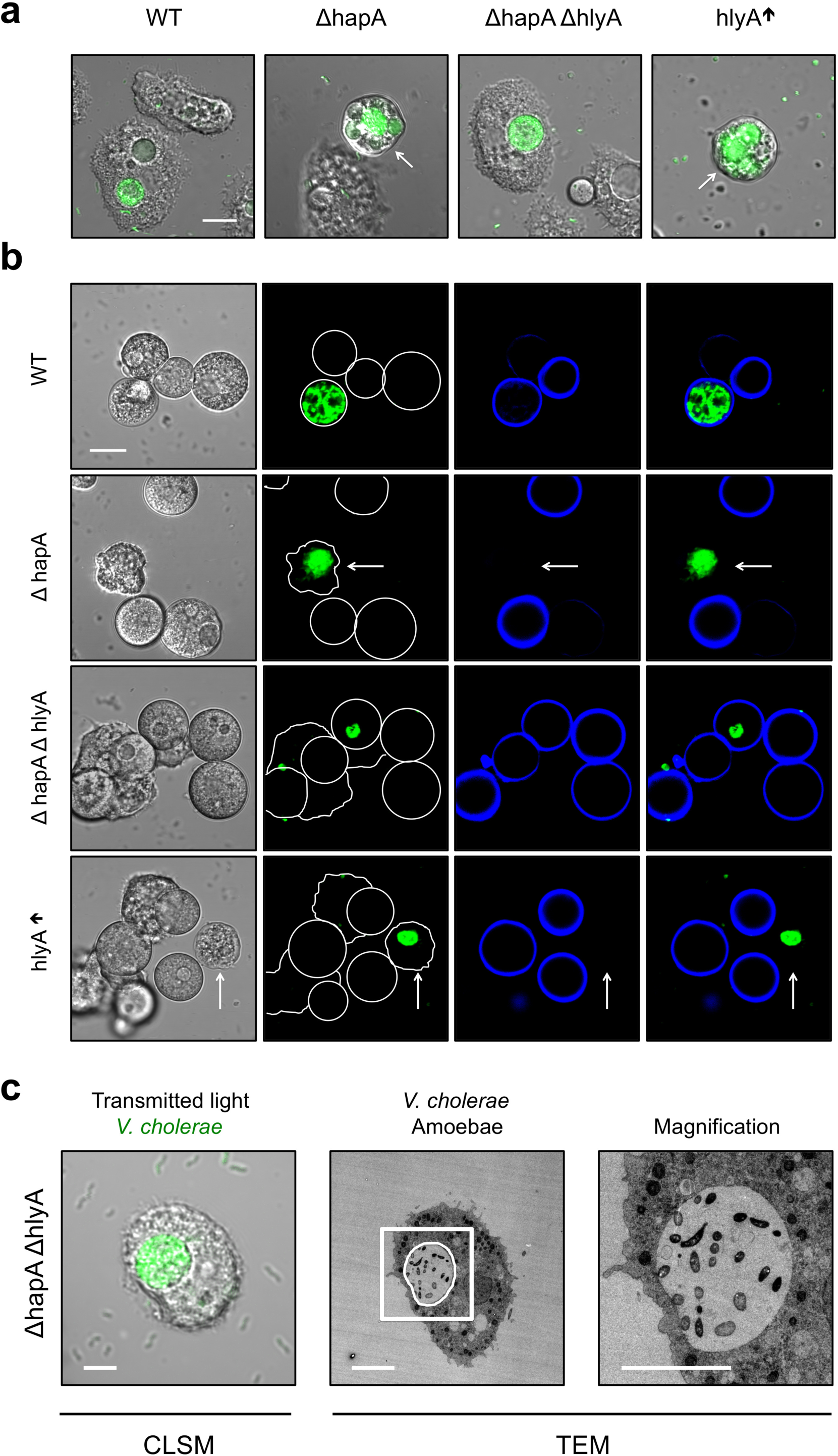
Amoebal intoxication in hemagglutinin protease deficient strains is caused by the pore-forming toxin hemolysin (HlyA) **(a)** Aberrant amoebal morphotypes are caused by hemolysin (HlyA). Deletion of the hemolysin gene (hlyA) in the protease-minus *V. cholerae* strain (ΔhapAΔhlyA) nullifies intoxication of the amoebal trophozoites upon infection, while the latter is elicited through the overexpression of *hlyA* in the WT background (hlyA↑). All of the bacterial strains were GFP-tagged. Merged images of the transmitted light channel and the green channel were taken at 20 hours p.p.c. Aberrantly shaped amoebae are indicated by white arrows. Scale bar: 10 μm **(b)** Absence of amoebal cellulose deposition is caused by hemolysin. *A. castellanii* cells were infected by the same *V. cholerae* strains as in panel (a) and, after staining with calcofluor, imaged at 30 hours p.p.c. White arrows depict colonized amoebae that are calcofluor-negative and aberrantly-shaped. Scale bar: 10 μm **(c)** CLEM image of an amoeba that is infected by the protease- and hemolysin-deficient *V. cholerae* strain.

To determine if HlyA impacts the amoebae through activity inside of the contractile vacuole instead of in the surrounding medium, we co-infected *A. castellanii* with WT *V. cholerae* (labeled with dsRed) and either a second WT strain as a control (Fig. S4a) or, alternatively, the *hapA*-minus strain or the HlyA-overexpression strain at a 1:1 ratio (all labeled with GFP; Fig S4b and c). We then imaged the amoebae and quantified the number of colonized ones that showed either a normal or aberrant morphology (Fig. S4). This experiment suggested that the secreted protease in the WT strain rescues the amoebae from intoxication by protease-minus and, therefore, HlyA-producing *V. cholerae* if both strains occurred within the same contractile vacuole (Fig. S4b and Table S2). This rescuing phenotype was not observed when the second strain was the HlyA-overproducing strain (Fig. S4c and Table S2). Amoebae containing mono-colonized contractile vacuoles showed the same phenotype as we had observed in the single strain infection experiments (Fig. S4 compared to Figs. 2 and 3). These *in vivo* observations are, therefore, fully consistent with the *in vitro* hemolysin activity described above (Fig. S2) and suggest that both enzymes are present and active inside the amoebal contractile vacuole.

To further support these findings and the interplay between the HapA protease and the HlyA hemolysin, we generated a gene encoding a translational fusion between HlyA and superfolder green fluorescent protein (sfGFP)^36^ and used this allele to replace the endogenous copy of *hlyA.* The resulting strain and a corresponding *hapA*-minus derivative (both also containing a constitutively expressed *dsRed* gene on their chromosome; Table S1) were used to confirm the functionality of the fusion construct, as judged by the hemolysis of blood cells (Fig. S5a). Next, we infected amoebae with these strains. Due to the constitutively produced red fluorescence we had a straightforwardly method for detecting colonized contractile vacuoles. We further witnessed that green fluorescence from the HlyA-sfGFP fusion was hardly detectable in the WT background, which contrasted with the protease-minus strain (Fig. S5b and quantified in Fig. S5c). Interestingly, the sfGFP signal was not uniformly distributed within the contractile vacuole but seemingly localized to the edge in a patchy manner (Fig. S5b), suggesting oligomerization and potential pore formation within the membrane.

### *V. cholerae* uses its lecithinase to lyse amoebal cysts

While the experiments described above highlighted the essential nature of the HapA protease for reducing the cytotoxicity of *V. cholerae* towards its amoebal host at early time points, it remained to be discovered how the pathogen eventually escaped from its amoebal host to return to the environment. We reasoned that the pathogen would need to disrupt the host’s plasma membrane. The plasma membrane of *A. castellanii* contains lecithin, which is a mixture of glycerophospholipids that includes a high percentage of phosphatidylethanolamine and phosphatidylcholine^37^. We therefore speculated that *V. cholerae* might use its lecithinase^25^ to disrupt this membrane. We generated a *lec*-minus strain and confirmed its impaired lecithinase activity on egg yolk plates (Fig. S6). We also complemented the *lec*-minus strain by placing a new copy of *lec* that was preceded by its native promoter onto the chromosome Δlec::lec). The same genetic constructs were also added into the WT background to generate a *lec* merodiploid strain of *V. cholerae* (WT::lec). All of the strains were tested for their lecithinase activity *in vitro* and behaved as expected (Fig. S6).

Next, we infected *A. castellanii* with this set of genetically engineered strains. We observed that while the *lec*-deficient strains were able to colonize the contractile vacuole and escape from it in the cyst stage, the bacteria were unable to ultimately lyse the amoebal host, as visualized through the exclusion of extracellular dextran (Fig. 4). The complemented strain had its cyst lysis capability restored, a phenotype that was enhanced for the *lec* merodiploid bacteria (Fig. 4). We therefore concluded that the lecithinase enzyme indeed targets and permeabilizes the plasma membrane of the host, thereby triggering the death of the cyst.

**Figure 4.**
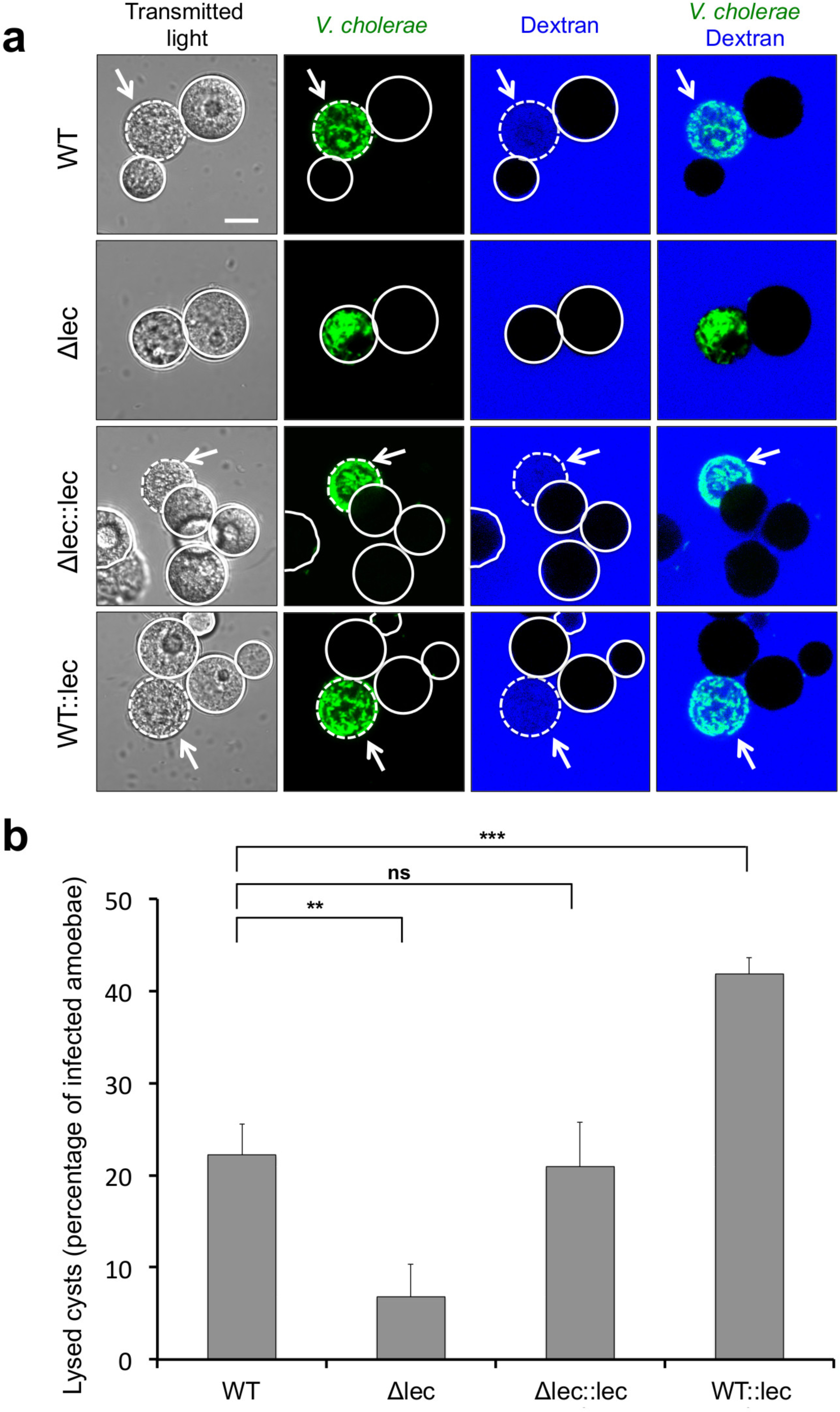
Cyst permeabilization by *V. cholerae* depends on its lecithinase enzyme. **(a)** Representative images and **(b)** quantification of lysed or non-lysed cysts that were colonized by WT, lecithinase-minus (Δlec), lecithinase-minus but complemented (△lec::lec), and *lec-* merodiploid *V. cholerae* strains (all GFP positive). The permeabilization phenotype was visualized through the infiltration of dextran from the surrounding medium in the white arrow-marked cysts, which did not occur for the Δlec strain. Scale bar: 10 μm. The graph in panel (b) represents average values from three independent biological replicates (± SD). Statistics are based on a one-way ANOVA with ***, *p* ≤ 0.001; **, *p* ≤ 0.01; ns, *p* > 0.05.

### *V. cholerae* escapes the lysed amoebal compartments through flagellum-based motility

As we observed that *V. cholerae* is highly motile within the amoebal host (Movie S2), we wondered whether this motility contributed to the pathogen’s intra-amoebal lifestyle and its escape from the succumbed host. We therefore generated non-motile mutants of *V. cholerae* by deleting either the gene that encodes the major flagellin subunit FlaA or the flagellar motor protein PomB (Table S1). While the first approach resulted in non-flagellated bacteria, the latter approach led to rotation-deficient but fully flagellated bacteria (Fig. 5A). The motility deficiency of the mutants was further confirmed in *in vitro* motility assays (Fig. S7). Next, we infected *A. castellanii* with these mutant strains. While both mutant strains still infected the amoebal contractile vacuole, the strains were more static within this niche (Fig. 5b and Movies S3 and S4). We therefore wondered whether motility would play a role in the escape of *V. cholerae* from the lysed contractile vacuole and lysed cysts. To test this idea, we infected amoebae with the WT and the *flaA*-minus strains, which we labeled with different fluorescent proteins, namely dsRed and GFP, respectively. We then took time-lapse movies over several hours to observe the colonizing bacteria of those amoebae that contained both strains within the same contractile vacuole (Fig. 5c and Movie S5). These experiments showed that, compared to the WT strain, the non-motile mutant was retained in the lysed contractile vacuole (Movie S6) and its escape back to the environment was severely impaired after the final lysis of the cyst (Movie S7). We concluded, therefore, that motility plays a major role in *V. cholerae*’s interaction with *A. castellanii.*

**Figure 5.**
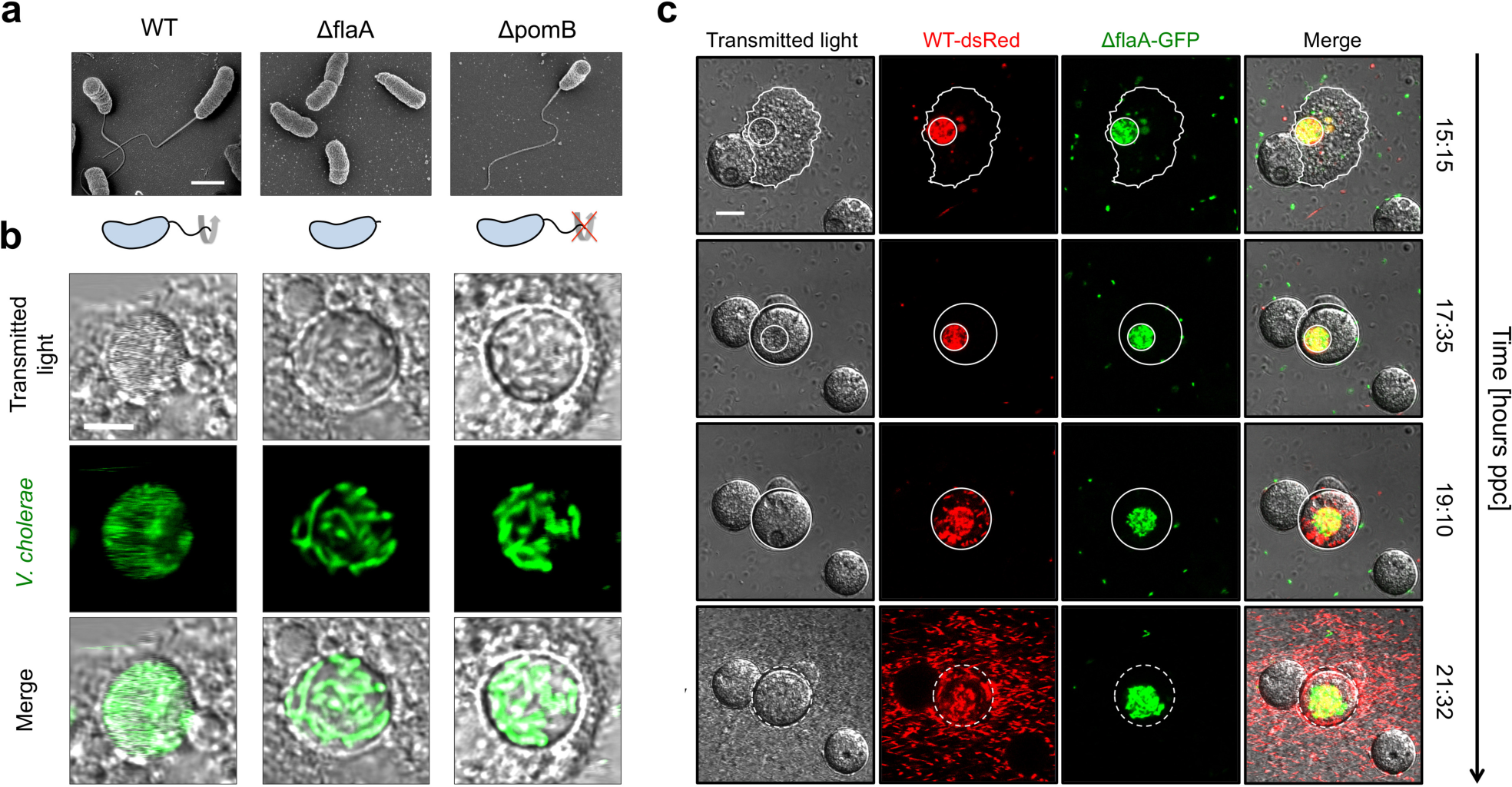
Efficient escape from the ruptured contractile vacuole and the lysed cysts requires flagellum-based motility. **(a)** Scanning electron micrographs of WT *V. cholerae* and its *flaA-* and *pomB*-deficient derivatives, which lack the major flagellin and cannot rotate their flagellum, respectively (as schematized below the images). Scale bar: 1 μm. **(b)** Close-up view of confocal scanning images of contractile vacuoles (transmitted light) that were colonized by each one of the three GFP-tagged strains shown in panel (a). The scanning speed was lowered for these experiments to visualize the fast movement of WT *V. cholerae* inside the contractile vacuole. See also Movies S2 to S4 for dynamics of intra-vacuolar WT, ΔflaA?, and ΔpomB bacteria. Scale bar: 5 μm. **(c)** Escaping from the lysed amoebal host and spreading requires flagellum-based motility. *A. castellanii* was infected with a 1:1 mixture of WT (dsRed-tagged) and non-motile *flaA*-minus (GFP-tagged) bacteria. Time-lapse microscopic imaging of a co-infected amoeba was started at 15 hours p.p.c. and followed for more than six hours (time is indicated on the right). Still images derived from the recorded movie (Movies S5) are depicted and show (from top to bottom) the colonized trophozoite before and after encystation, after rupture of the contractile vacuole, and after cyst lysis. Scale bar: 10 μm.

## Discussion

*V. cholerae*, the bacterial agent responsible for cholera, still poses a global threat to human health. However, apart from its chitin-induced phenotypes, which include chitin catabolism, inter-bacterial competition, and horizontal gene transfer^38^ (reviewed by^39-42^), we know very little about its environmental lifestyle and its potential interactions with non-human hosts. Such ancient host-pathogen interactions are, however, often considered as evolutionary precursors to modern interactions that occur between bacteria and their human hosts^43^. Moreover, aquatic predators are recognized for their contribution to pathogen emergence due to the selection pressure they exert on their bacterial prey^44^. Here, we examined the molecular mechanisms that *V. cholerae* uses to interact with the aquatic amoeba *A. castellanii* and to maintain a favorable replication niche within the amoebal osmoregulatory organelle. Based on the molecular checkpoints that were deciphered in the current study, we expanded our model of the pathogen’s intra-amoebal lifecycle (Fig. 6). Specifically, we showed the importance of several extracellular enzymes in this host-pathogen interaction. The production of the HapA protease, which cleaves the pore-forming hemolysin toxin, proved essential for avoiding premature intoxication of the amoebal host (Fig. 6a). In contrast to intracellular pathogens, such as *Listeria monocytogenes* or *Shigella flexneri*, that use pore-forming toxins to escape from acidic vacuolar compartments to reach the host cell’s cytosol, the situation described herein is very different. In this case, the host-pathogen interaction relies on *V. cholerae* residing in a non-digestive vacuole in which it can readily replicate^30^. This nondigestive contractile vacuole is an essential osmoregulatory organelle of the amoeba meaning that HlyA-mediated rupture of the vacuolar membrane would, therefore, release *V. cholerae* into the cytosol, though, at the expense of rapid host cell death. We speculated, therefore, that this HapA-mediated disintegration of HlyA might have evolved to allow *V. cholerae* to maximize its growth output within this intra-amoebal replication niche by avoiding the premature death of its host. Notably, the HapA protease is considered a minor virulence factor that contributes to disease outcome in animal models of cholera^15,45^, along with HlyA itself, as described above. In this study, we did not observe any obvious phenotype for *V. cholerae* strains lacking HlyA with respect to their ability to first colonize and then escape the amoebal contractile vacuole. However, at this point we cannot exclude the contribution of HlyA as well as other, potentially redundant, enzymes to the other discussed phenotype in which the *V. cholerae* pathogen escapes from the *A. castellanii* phagosomal pathway^30^. Alternatively, HlyA might play a role in another context in the pathogen’s environmental lifestyle. It was shown that HlyA causes developmental delays and lethality in the nematode *Caenorhabditis elegans* through intestinal cytopathic changes^46^, and similar effect might occur with other hosts in the pathogen’s aquatic reservoirs. Notably, both enzymes (HapA and HlyA) are widely distributed among pathogenic *Vibrios*^17^, which reinforces the notion that they play important roles in environmental settings.

**Figure 6.**
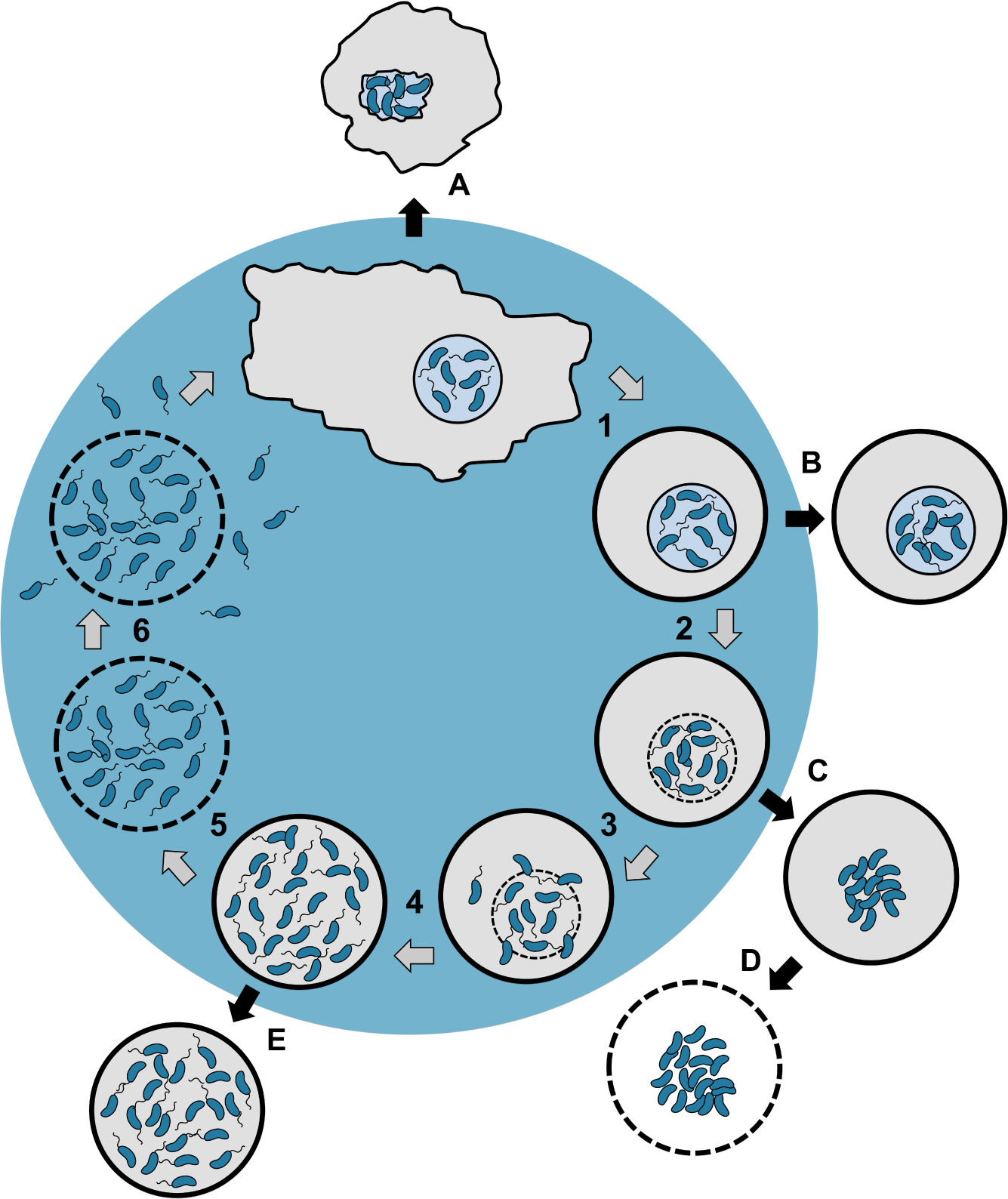
Model depicting the herein-described molecular checkpoints of *V. cholerae’s* amoebal infection cycle. After phagocytosis, *V. cholerae* cells colonize the contractile vacuole of *A. castellanii* trophozoites. The bacteria remain within this replication niche upon amoebal encystation (1). After intra-vacuolar growth, rupture of the vacuole occurs in a *Vibrio* polysaccharide-dependent manner^30^ (2), which releases motile *V. cholerae* into the cyst’s cytosol (3) where the bacteria proliferate further (4). Lecithinase-mediated membrane permeabilization (5) eventually leads to the lysis of the cyst and the quick release of the motile bacteria (6), which can undergo another round of infection. Defined bacterial mutants are blocked at different checkpoints and, therefore, impaired in the progression of the infection cycle: **A**, protease-deficient and therefore hemolysis-overactive strain (ΔhapA); **B**, *Vibrio* polysaccharide (VPS)-deficient strain ΔvpsA^30^); **C** and **D**, non-motile strains ΔflaA and ΔpomB); and **E**, lecithinase-minus strain (Δlec).

While the production of the HapA protease protects the contractile vacuole from premature lysis, such lysis still occurs when the amoeba has undergone encystation and the contractile vacuole is fully occupied by the bacteria. We have previously shown that the rupture of the vacuolar membrane is dependent on the production of the *Vibrio* extracellular polysaccharide (VPS; Fig. 6b), and we hypothesized that this might be due to a VPS-dependent agglutination effect^30^. Consistent with this idea is a recent study on the importance of the osmotic pressure generated by the hydrogel-like VPS matrix of *V. cholerae^47^.* These authors suggested that the capacity of *V. cholerae* to escape the amoebal contractile vacuole, which we had previously reported^30^, could indeed be related to the osmotic-pressure response of the VPS biofilm matrix^47^.

After *V. cholerae* escaped from the vacuole and upon further growth within the cytosol of the amoebal cyst, another extracellular enzyme, the lecithinase/phospholipase, was required to ultimately kill the host and return the bacteria to the environment (Fig. 6e). This lecithinase is encoded in close proximity to the hemolysin gene, a region that was previously speculated to represent a pathogenicity island^23^. Unfortunately, apart from the absence of a phenotype for the *lec*-minus mutant in rabbit-ligated ileal loops, a model of cholera that primarily aims at judging cholera toxin-mediated fluid accumulation^25^, this enzyme had not been extensively studied *in vivo* (for current 7^th^ pandemic O1 El Tor strains). However, a recent study based on an activity-based protein profiling method showed that the lecithinase/phospholipase is active in cecal fluids of infected infant rabbits^48^. Further studies are therefore required to conclusively determine whether this enzyme should also be considered as a minor virulence factor for animals, as we demonstrated here for amoebae.

Lastly, we showed that bacterial motility is required to efficiently escape from the lysed contractile vacuole and the succumbed host (Fig. 6e), which, ultimately, allows the bacteria to return to the aquatic environment. One could speculate that the lysed cyst might generate a nutrient gradient that attracts novel predators, a disadvantageous scenario for any prey. Indeed, we frequently observed that dead cysts were readily ingested by feeding trophozoites, which could counter-select for non-motile mutants that cannot escape from the cellular debris. Likewise, motility has also been demonstrated to be of importance for virulence in animal models. Indeed, non-motile mutants exerted less fluid accumulation in rabbit ileal loops^45^ and were severely attenuated for the colonization of infant mouse small intestines^6^.

These correlations between the contribution to virulence in animal models and in the environmental host-pathogen interaction described herein therefore support the “coincidental evolution hypothesis”. This hypothesis suggests that “virulence factors result from adaptation to other ecological niches” and, in particular, from “selective pressure exerted by protozoan predator”^49^.

## Methods

### Bacterial strains and growth conditions

All bacterial strains used in this study are listed in Table S1. *V. cholerae* strains are derivatives of the 7^th^ pandemic O1 El Tor strain A1552^50^. Bacteria were grown on LB plates (1.5% agar) or under shaking conditions in liquid LB medium. Antibiotics were supplemented when needed for genetic engineering and selection at the following concentrations: 100 μg/ml of ampicillin; 50 μg/ml of gentamicin, and 75 μg/ml of kanamycin. Amoebal infection media were free of antibiotics.

For counter-selection of site-directly modified phenylalanyl-tRNA synthetase (*pheS*\*)-carrying strains (Trans2 method; see below) 4-chloro-phenylalanine (cPhe; C6506, Sigma-Aldrich, Buchs, Switzerland) was added to the medium before autoclaving. Optimization experiments showed that a concentration of 20 mM of cPhe was best for *V. cholerae* counter selection.

### Genetic engineering of bacterial strains

Genetic engineering of strains to delete/insert genes or to add gene fusions onto the *V. cholerae* chromosomes was performed using the previously described TransFLP method^51-54^. An alternative protocol, based on two rounds of transformation, was established in this study (named Trans2) to generate complemented strains or overexpression constructs. This method was adapted from Gurung and colleagues^55^ to work in *V. cholerae* and is based on the counter-selectablility of bacteria carrying a mutated version of the α subunit of phenylalanyl- tRNA synthetase in cPhe-containing medium. The gene encoding the phenylalanyl-tRNA synthetase of *V. cholerae* (α subunit; *pheS* or VC1219) was integrated into plasmid pBR-FRT-Kan-FRT2^56^ to generate the plasmid pBR-FKpheSF (carrying the aminoglycoside 3’- phosphotransferase-encoding gene [*aph*(3’)] and *pheS* flanked by FRT sites). This and subsequent plasmids served as templates for inverse PCR to generate the site-directly mutated *pheS* in the pBR-FKpheSF[A294G] and pBR-FKpheSF[A294G/T251A] plasmids (renamed for simplicity as pFRT-aph-pheS&). This double-mutant template was generated based on a study in *Escherichia coli* that showed improved counter-selectablility of this version of pheS (*pheS*\*) on cPhe-containing agar plates^57^. In contrast to a previous study^55^, the *pheS*\* allele used in this study (pheS[A294G/T251A]) was placed downstream of the constitutively expressed *aph*(3’) to ensure expression independent of the subsequent insertion site on the chromosome. The *aph*(3’)-*pheS*\* construct was PCR amplified using the pFRT-aph-pheS* plasmid as a template and fused by overlapping PCR with fragments carrying parts of the 5’- and 3’-region of the *lacZ* gene. The resulting PCR fragment (*lacZ’-aph*(3’)-*pheS*-‘lacZ*) was used as a transforming material for chitin-induced naturally competent *V. cholerae* cells^52^. Transformants were selected on kanamycin-containing LB agar plates. The growth impairment of the transformants due to the presence of *pheS*\* was verified on cPhe-containing agar plates followed by verification through Sanger sequencing. These strains underwent a second round of transformation on chitin (therefore the name Trans2). The second-round transforming material was devoid of any resistance gene and carried the desired genetic construct (generated by PCR amplification) flanked by the 5’- and 3’-parts of the *lacZ* gene. Screening for the integration of this construct was done using cPhe-containing agar plates. Transformants that grew were tested by colony PCR for the replacement of the *aph*(3’)-*pheS*\* genes and, ultimately, confirmed for mutation-free and marker- and scar-less integration using Sanger sequencing.

Variants of the miniTn7 transposon carrying constitutively expressed *gfp* or an optimized version of *dsRed* (*dsRed.T3*[*DNT*]^38,58,59^) were stably integrated into the *V. cholerae* chromosome through triparental mating^60^.

### Gene expression analysis based on qRT-PCR

The relative gene expression was determined using quantitative reverse transcription PCR (qRT-PCR)-based transcript scoring in *V. cholerae* as previously described^61^. Data are based on three biologically independent experiments (± standard deviation [SD]).

### Amoebal infections

The amoeba *Acanthamoeba castellanii* (ATCC strain 30010) served as the host in all of the infection experiments. Supplemented peptone-yeast-glucose (PYG) medium (ATCC medium 712) was used to propagate uninfected amoebae and half-concentrated defined artificial seawater (0.5× DASW; buffered with 50 mM HEPES)^62^ was used as the infection medium.

Infection experiments were performed according to a previous study^30^. Briefly, amoebae were diluted with fresh PYG at a concentration of 1 × 10^5^ amoebae/ml and seeded into a μ-Dish (low wall 35 mm ibidiTreat devices; 80136-IBI, Vitaris, Baar, Switzerland). After three hours of static incubation at 24°C, adherent amoebae were washed three times with the infection medium. Exponentially growing bacteria were likewise extensively washed in infection medium and added to the amoebae at a multiplicity of infection (MOI) of 1000. Cocultures were incubated statically at 24°C for the indicated time post-primary contact (p.p.c.).

### Confocal laser scanning microscopy-based techniques

Confocal lasers scanning microscope (CLSM) imaging was done using a Zeiss LSM 700 inverted microscope (Zeiss, Feldbach, Switzerland). To label the endosomal pathway of the amoebae^63^, Alexa Fluor 647-labeled dextran (MW 10,000; D22914, Thermo Fisher Scientific, Waltham, Massachusetts, USA) was added to the infection medium at a final concentration of 100 μg/ml. The fluorescent stain calcofluor white was supplemented wherever indicated at a final concentration of 0.2%, as this stain is known to bind to cellulose (18909-100ML-F, Sigma-Aldrich Chemie GmbH, Buchs, Switzerland). Fluorescent signals were quantified using the open-source imaging software ImageJ^64^ (NIH, Bethesda, MD, USA). Bacterial motility inside the amoebae (see Movies S2, S3, and S4) was assessed using short frame intervals and/or a reduced scanning speed.

### Scanning electron microscopy

For scanning electron microscopy (SEM) imaging, a 10 mm in diameter silicon wafer was coated with 100 μg/ml of poly-D-lysine hydrobromide (P6407, Sigma-Aldrich, Buchs, Switzerland) for two hours at room temperature, then washed three times with bi-distillated water. Exponentially grown bacteria were washed with PBS and allowed to settle for 20 minutes at room temperature onto the silicon wafer. The supernatant was removed, and a 0.1 M phosphate buffer solution of 1.25% glutaraldehyde and 1% tannic acid was gently added. After one hour, the fixative was removed and replaced by cacodylate buffer. This was transferred to a 1% solution of osmium tetroxide in cacodylate buffer for 30 minutes, washed in water, and then dehydrated through a series of increasing concentrations of alcohol. The sample was finally dried at the critical point of carbon dioxide and coated with a 2 nm-thick layer of osmium metal using an osmium plasma coater. Images were collected in an SEM (Merlin, Zeiss NTS) at 1.8 kV electron beam tension.

### Correlative light and electron microscopy

For the correlative light and electron microscopy (CLEM), 18 mm glass coverslips were used. These were coated with a thin (10 nm) layer of carbon onto which an alphanumeric grid had been outlined using a thicker, 20 nm, carbon layer. The coverslips were incubated with poly-D-lysine hydrobromide (P6407, Sigma-Aldrich, Buchs, Switzerland) at a concentration of 100 μg/ml for 30 minutes at room temperature and then washed three times with PBS. At 20 hours p.p.c., colonized amoebae were detached from the tissue culture flask using a cell scraper, plated onto coated coverslips, and allowed to adhere for 30 minutes at room temperature. The supernatant was removed, and a buffered aldehyde solution of 1% glutaraldehyde and 2% paraformaldehyde was gently added. After one hour at room temperature, these were washed in cacodylate buffer. The corresponding confocal images were acquired. The samples were then post-fixed and stained with 1% osmium tetroxide followed by 1% uranyl acetate. After dehydration through increasing concentrations of alcohol, the coverslips were embedded in EPON resin and polymerized at 65°C for 48 hours. Cells of interest were located using the alphanumeric grid that remained on the resin surface after the grid had been removed. Blocks were trimmed around the cell and between 200 to 300 serial thin sections, at a thickness of 50 nm, were cut from the entire structure using a diamond knife and ultra-microtome. Sections were collected onto copper slot grids, carrying a formvar support film, and then further stained with uranyl acetate and lead citrate. Serial images of the cell were taken with a transmission electron microscopy (TEM; Tecnai Spirit, Fischer Scientific) at a voltage of 80 kV.

To generate a 3D model of the colonized contractile vacuole, serial TEM images were first aligned manually using image software (Photoshop, Adobe). Each structure was then segmented using TrakEM2^65^ operating in the FIJI software^66^ (www.fiji.sc). The 3D models were then exported into the Blender software (www.blender.org) for smoothing and rendering into a final image or movie.

### Determination of enzymatic activities

Semi-quantitative assays in specific media were used to determine enzymatic activities. To perform these assays, 5μl of the respective overnight bacterial cultures were spotted onto the following three types of agar plates. (i) Milk agar was used to determine protease activity due to the hydrolysis of casein. The modified recipe used in this study contained 0.5% tryptone, 0.25% yeast extract, 0.1% dextrose, 1% skim milk powder, and 1.25% agar. The bacteria were incubated on these plates for 24 hours at 30°C. (ii) Trypticase Soy Agar II with 5% sheep blood (BD, Heidelberg, Germany) was used to test for hemolysis (24 hours at 30°C). (iii) Egg yolk agar plates (Hardy Diagnostic, Santa Maria, CA, USA) were used to evaluate lecithinase activity (72 hours at 30°C). Lecithinase activity was quantified after taking pictures with a tabletop scanner and analyzing these images using ImageJ. Values are given according to the formula (area of bacteria + area of precipitate) – (area of the bacteria) = area of precipitate [cm^2^].

### Statistical analysis

One-way ANOVA was performed using GraphPad Prism version 7 for Mac (GraphPad Software, San Diego, CA, USA) followed by Tukey’s post hoc test for multiple comparisons. Statistical analyses were performed on three biologically independent experiments. * *p* ≤ 0. 05; **, *p* ≤ 0.01; ***, *p* ≤ 0.001; ****, *p* ≤ 0.0001; ns (not significant), *p* > 0.05. The amoebal population counted was each time 6,000 (derived from three biologically independent experiments each with n=2,000).

### Data availability

All data supporting the findings of this study are available from the corresponding author upon request.

## Acknowledgements

The authors thank members of the Blokesch group at EPFL and the Geneva/Lausanne amoeba club for fruitful discussions. We also acknowledge former members of the Blokesch lab for the contribution of some *V. cholerae* strains. This work was supported by EPFL intramural funding and a Starting Grant from the European Research Council (ERC; 309064-VIR4ENV) to MB. M.B. is a Howard Hughes Medical Institute (HHMI) International Research Scholar.

## Author contributions

C.V. and M.B. designed the study; C.V., G.K. and M.B. planned the experiments; C.V., S.S., C.S., T.S., S.R., C.M., and M.B. performed experiments; C.V. and M.B. analyzed the data; C.V. and M.B. wrote the manuscript. All authors approved the manuscript.

## Additional information

Supplementary information is available online. Correspondence and requests for materials should be addressed to M.B.

## Competing interests

The authors declare no competing financial interests.

**Figure legends - supplementary figures**

**Figure S1**. **Aberrant amoebal morphology caused by hemagglutinin protease (HapA)**-deficient *V. cholerae*. Additional confocal images of GFP-tagged bacteria and amoebae as in Fig. 2a. Shown are merged images of the transmitted light channel and the green channel. Bacterial strains depicted: WT, ΔhapA, and the complemented mutant (ΔhapA::hapA). White arrows depict aberrant morphotypes.

**Figure S2. Characterization of genetically engineered *V. cholerae* strains. Diverse *V. cholerae*** strains were tested *in vitro* **(a)** for hemolytic activity on blood agar plates and for protease activity on milk agar plates or **(b)** for expression of representative genes by qRT-PCR. The strains tested in panel (a) are WT, ΔhapA, ΔhapA-complemented (ΔhapA::hapA), the protease and hemolysin double mutant without *hlyA* complementation (ΔhapAΔhlyA) and with *hlyA* complementation (ΔhapAΔhlyA::hlyA), and a *hlyA*-overexpression strain (hlyA↑). The last two rows show 1:1 mixtures of both indicated strains. **(b)** Relative gene expression confirming proficient complementation or overexpression of *hapA* and *hlyA*. Transcript levels were scored by qRT-PCR and normalized to *gyrA* (shown as relative expression values on the 7-axis). The gene encoding the quorum-sensing master regulator HapR (*hapR*) serves as negative control (no difference in its expression in any of the tested strains). Strains tested: 1, WT; 2, ΔhapA; 3, ΔhapA::hapA; 4, ΔhlyA; 5, ΔhlyA::hlyA; 6, ΔhapAΔhlyA; 7, ΔhapAΔhlyA::hlyA; 8, hlyA↑. Data are averages from three independent biological replicates (± SD).

**Figure S3. The hemolysin of *V. cholerae* intoxicates *A. castellanii.* (a), (b)** Additional confocal micrographs of amoebae infected with GFP-tagged ΔhapAΔhlyA and hlyA↑ strains of *V. cholerae.* Details as in Fig. 3a. Shown are in panel (a) the transmitted light channel, the green channel, and the green channel and in panel (b) merged channel images only. Scale bars for (a) and (b): 10 μm. The white arrows show aberrant morphotypes. **(c)**, **(d)** Quantification of normal or aberrant morphologies and deposition of cellulose by amoebae infected with hemolysin-overactive *V. cholerae* strains. Details as in Fig. 2b and e. Values for WT, ΔhapA, and ΔhapA::hapA are the same as in Fig. 2 to compare the values to the ΔhapAΔhlyA and hlyA↑ strains. Statistics are based on a one-way ANOVA with **** *p* ≤ 0.0001; ns, *p* > 0.05. **(e)** Malformation of the contractile vacuole in amoebae infected by protease-deficient and, therefore, hemolysin-overactive *V. cholerae* strains. Additional CLEM images as in Fig. 3c show the malformation of the contractile vacuole in *hapA-* deficient strains (ΔhapA) compared to WT or *hapA-* and *hlyA*-deficient strains (ΔhapAΔhlyA).

**Figure S4. Intra-vacuolar activities of HapA and HlyA**. Amoebae infected by a 1:1 mixture of the two strains were imaged by confocal microscopy at 20 hours p.p.c. The WT is tagged with dsRed in all images, while the GFP-tagged strain corresponds to **(a)** a second WT, **(b)** ΔhapA, **(c)** and hlyA↑. The micrographs depict amoebae colonized (I) by the dsRed-tagged strain, (II) the GFP-tagged strain, (III) or both strains. For each lane, the transmitted light channel, the red channel, the green channel, and a merged image of all three channels are displayed. Quantifications of these experiments from three independent biological replicates are given in Table S2. #Micrograph shows a single event observed out of 6,000 amoebae counted.

**Figure S5. GFP-tagged hemolysin can be visualized in amoebae infected with hemagglutinin protease-deficient *V. cholerae.* (a)** *In vitro* characterization of engineered bacterial strains. *V. cholerae* strains containing constitutively expressed *dsRed* in the WT, ΔhapA, or ΔhapAΔhlyA background without or with an allele encoding for superfolder GFP fused to hemolysin (*hlyA-sfGFP*) were tested for hemolysin activity on blood agar plates (scored below the images). **(b)** The HlyA-sfGFP fusion protein is visible in the contractile vacuole of amoebae infected by the hemagglutinin protease-deficient *V. cholerae* strain. Depicted are confocal micrographs of the transmitted light channel, the red channel, the green channel, and the merged fluorescent channels (from left to right). Scale bar: 10 μm. **(c)** Quantification of HlyA-sfGFP-derived green fluorescence in the WT and ΔhapA background. The green fluorescent signal was normalized against the red fluorescent signal (derived from constitutively expressed *dsRed*) and given as arbitrary units (AU; indicated on the 7-axis). The WT-dsRed strain that does not carry the *hlyA-sfGFP* allele served as the negative control to correct for background green fluorescence. The data was pooled from three independent experiments and each dot represents a singly analyzed, colonized amoeba with the lines showing the average values. Statistics are based on one-way ANOVA. *, *p* ≤ 0.05; ****, *p* ≤ 0.0001.

**Figure S6. Quantification of *in vitro* lecithinase activity exerted by diverse *V. cholerae* strains. (a)** *V. cholerae* strains were tested for lecithinase activity on egg yolk agar plates. Lecithinase degrades lecithin that is present in egg yolk, thereby producing an insoluble and opaque precipitate around the colonies. **(b)** Quantification of the area of precipitation. Bacterial strains tested were WT, Δlec, Δlec-complemented (Δlec::lec), and the *lec-* merodiploid strain (WT::lec). The graph shows the averages of three independent biological experiments (± SD). Statistics are based on one-way ANOVA. **, *p* ≤ 0.01; ****, *p* ≤ 0.0001; ns,*p* > 0.05.

**Figure S7. Confirmation of *in vitro* motility of genetically engineered *V. cholerae* strains.**

The *V. cholerae* WT, ΔflaA and ΔpomB strains (without or with constitutively expressed *gfp* or *dsRed*, as indicated) were tested for flagellum-based motility on soft agar plates. **(a)** Representative images. **(b)** Data from three independent biological replicates were quantified based on the swarming diameter.

**Table S1.** Bacterial strains and plasmids used in this study.

**Table S2.** Quantification of bacterial hemolysin-triggered aberrant amoebal morphologies.

**Movie S1:** 3D reconstruction of a *V. cholerae*-colonized amoebal contractile vacuole.

**Movie S2:** Intra-vacuolar dynamics of WT *V. cholerae.*

**Movie S3:** Intra-vacuolar dynamics of ΔflaA *V. cholerae.*

**Movie S4:** Intra-vacuolar dynamics of ΔpomB *V. cholerae.*

**Movie S5:** Time-lapse confocal microscopy movie of a colonized contractile vacuole cohousing WT (dsRed-tagged) and ΔflaA (GFP-tagged) bacteria. See snapshots in Fig. 5c.

**Movie S6:** Short section of Movie S5 at a slower speed showing the rupture of the contractile vacuole.

**Movie S7:** Short section of Movie S5 at a slower speed showing the cyst lysis and the spread of WT bacteria.

## References

1. Faruque, S. M., Albert, M. J. & Mekalanos, J. J. Epidemiology, genetics, and ecology of toxigenic Vibrio cholerae. Microbiol. Mol. Biol. Rev. 62, 1301–1314, (1998).

2. Clemens, J. D., Nair, G. B., Ahmed, T., Qadri, F. & Holmgren, J. Cholera. Lancet 390, 1539–1549, (2017).

3. Ali, M., Nelson, A. R., Lopez, A. L. & Sack, D. A. Updated global burden of cholera in endemic countries. PLoSNegl. Trop. Dis. 9, e0003832, (2015).

4. WHO. World Health Organization: Cholera, 2015. Weekly Epidemiological Record 91, 433–440, (2016).

5. Chen, L. et al. VFDB: a reference database for bacterial virulence factors. Nucleic Acids Res. 33, D325–328, (2005).

6. Lee, S. H., Butler, S. M. & Camilli, A. Selection for *in vivo* regulators of bacterial virulence. Proc. Natl. Acad. Sci. USA 98, 6889–6894, (2001).

7. Flluner, K. J. in 8Microbial Pathogenesis and the Intestinal Epithelial Cell Ch. 26, 481–502 (ASM Press, 2003).

8. Zampini, M. et al. Vibrio cholerae persistence in aquatic environments and colonization of intestinal cells: involvement of a common adhesion mechanism. FEMSMicrobiol. Lett. 244, 267–273, (2005).

9. Kirn, T. J., Jude, B. A. & Taylor, R. K. A colonization factor links *Vibrio cholerae* environmental survival and human infection. Nature 438, 863–866, (2005).

10. Booth, B. A., Boesman-Finkelstein, M. & Finkelstein, R. A. *Vibrio cholerae* soluble hemagglutinin/protease is a metalloenzyme. Infect. Immun. 42, 639–644, (1983).

11. Burnet, F. M. & Stone, J. D. Desquamation of intestinal epithelium in vitro by *V. cholerae* filtrates; characterization of mucinase and tissue disintegrating enzymes. Aust. J. Exp. Biol. Med. Sci. 25, 219–226, (1947).

12. Finkelstein, R. A., Boesman-Finkelstein, M. & Holt, P. *Vibrio cholerae* hemagglutinin/lectin/protease hydrolyzes fibronectin and ovomucin: F.M. Burnet revisited. Proc. Natl. Acad. Sci. USA 80, 1092–1095, (1983).

13. Wu, Z., Milton, D., Nybom, P., Sjo, A. & Magnusson, K. E. *Vibrio cholerae* hemagglutinin/protease (HA/protease) causes morphological changes in cultured epithelial cells and perturbs their paracellular barrier function. Microb. Pathog. 21, 111–123, (1996).

14. Mel, S. F., Fullner, K. J., Wimer-Mackin, S., Lencer, W. I. & Mekalanos, J. J. Association of protease activity in *Vibrio cholerae* vaccine strains with decreases in transcellular epithelial resistance of polarized T84 intestinal epithelial cells. Infect. Immun. 68, 6487–6492, (2000).

15. Benitez, J. A. et al. Preliminary assessment of the safety and immunogenicity of a new CTXPhi-negative, hemagglutinin/protease-defective El Tor strain as a cholera vaccine candidate. Infect. Immun. 67, 539–545, (1999).

16. Finkelstein, R. A., Boesman-Finkelstein, M., Chang, Y. & Häse, C. C. *Vibrio cholerae* hemagglutinin/protease, colonial variation, virulence, and detachment. Infect. Immun. 60, 472–478, (1992).

17. Zhang, X. H. & Austin, B. Haemolysins in *Vibrio* species. J. Appl. Microbiol. 98, 1011–1019, (2005).

18. Menzl, K., Maier, E., Chakraborty, T. & Benz, R. HlyA hemolysin of *Vibrio cholerae* O1 biotype E1 Tor. Identification of the hemolytic complex and evidence for the formation of anion-selective ion-permeable channels. European journal of biochemistry 240, 646–654, (1996).

19. Ichinose, Y. et al. Enterotoxicity of El Tor-like hemolysin of non-O1 *Vibrio cholerae*. Infect. Immun. 55, 1090–1093, (1987).

20. Alm, R. A., Mayrhofer, G., Kotlarski, I. & Manning, P. A. Amino-terminal domain of the El Tor haemolysin of *Vibrio cholerae* O1 is expressed in classical strains and is cytotoxic. Vaccine 9, 588–594, (1991).

21. Olivier, V., Haines, G. K., 3rd, Tan, Y. & Satchell, K. J. Hemolysin and the multifunctional autoprocessing RTX toxin are virulence factors during intestinal infection of mice with *Vibrio cholerae* El Tor O1 strains. Infect. Immun. 75, 5035–5042, (2007).

22. Olivier, V., Salzman, N. H. & Satchell, K. J. Prolonged colonization of mice by *Vibrio cholerae* El Tor O1 depends on accessory toxins. Infect. Immun. 75, 5043–5051, (2007).

23. Ogierman, M. A. et al. Characterization of the *Vibrio cholerae* El Tor lipase operon *lipAB* and a protease gene downstream of the *hly* region. J. Bacteriol. 179, 7072–7080, (1997).

24. Felsenfeld, O. The Lecithinase Activity of Vibrio comma and the El Tor Vibrio. J. Bacteriol. 48, 155–157, (1944).

25. Fiore, A. E., Michalski, J. M., Russell, R. G., Sears, C. L. & Kaper, J. B. Cloning, characterization, and chromosomal mapping of a phospholipase (lecithinase) produced by *Vibrio cholerae*. Infect. Immun. 65, 3112–3117, (1997).

26. Vezzulli, L., Guzman, C. A., Colwell, R. R. & Pruzzo, C. Dual role colonization factors connecting *Vibrio cholerae’s* lifestyles in human and aquatic environments open new perspectives for combating infectious diseases. Curr. Opin. Biotechnol. 19, 254–259, (2008).

27. Halpern, M. Novel insights into Haemagglutinin Protease (HAP) gene regulation in *Vibrio cholerae*. Mol. Ecol. 19, 4108–4112, (2010).

28. Colwell, R. R. Global climate and infectious disease: the cholera paradigm. Science 274, 2025–2031, (1996).

29. Abd, H., Saeed, A., Weintraub, A., Nair, G. B. & Sandström, G. *Vibrio cholerae* O1 strains are facultative intracellular bacteria, able to survive and multiply symbiotically inside the aquatic free-living amoeba *Acanthamoeba castellanii*. FEMS Microbiol. Ecol. 60, 33–39, (2007).

30. Van der Henst, C., Scrignari, T., Maclachlan, C. & Blokesch, M. An intracellular replication niche for *Vibrio cholerae* in the amoeba *Acanthamoeba castellanii*. ISME J. 10, 897–910, (2016).

31. West, M. J. Stereological methods for estimating the total number of neurons and synapses: issues of precision and bias. Trends Neurosci. 22, 51–61, (1999).

32. Pal, R. A. The Osmoregulatory System of the Amoeba, *Acanthamoeba Castellanii*. J. Exp. Biol. 57, 55, (1972).

33. Lorenzo-Morales, J. et al. Glycogen phosphorylase in *Acanthamoeba* spp.: determining the role of the enzyme during the encystment process using RNA interference. Eukaryot. Cell 7, 509–517, (2008).

34. Dal Peraro, M. & van der Goot, F. G. Pore-forming toxins: ancient, but never really out of fashion. Nat. Rev. Microbiol. 14, 77–92, (2016).

35. Tsou, A. M. & Zhu, J. Quorum sensing negatively regulates hemolysin transcriptionally and posttranslationally in *Vibrio cholerae*. Infect. Immun. 78, 461–467, (2010).

36. Pedelacq, J. D., Cabantous, S., Tran, T., Terwilliger, T. C. & Waldo, G. S. Engineering and characterization of a superfolder green fluorescent protein. Nat. Biotechnol. 24, 79–88, (2006).

37. Ulsamer, A. G., Wright, P. L., Wetzel, M. G. & Korn, E. D. Plasma and phagosome membranes of *Acanthamoeba castellanii*. The Journal of cell biology 51, 193–215, (1971).

38. Borgeaud, S., Metzger, L. C., Scrignari, T. & Blokesch, M. The type VI secretion system of *Vibrio cholerae* fosters horizontal gene transfer. Science 347, 63–67, (2015).

39. Hunt, D. E., Gevers, D., Vahora, N. M. & Polz, M. F. Conservation of the chitin utilization pathway in the *Vibrionaceae*. Appl. Environ. Microbiol. 74, 44–51, (2008).

40. Matthey, N. & Blokesch, M. The DNA-Uptake Process of Naturally Competent *Vibrio cholerae*. Trends Microbiol. 24, 98–110, (2016).

41. Metzger, L. C. & Blokesch, M. Regulation of competence-mediated horizontal gene transfer in the natural habitat of *Vibrio cholerae*. Curr. Opin. Microbiol. 30, 1–7, (2016).

42. Veening, J. W. & Blokesch, M. Interbacterial predation as a strategy for DNA acquisition in naturally competent bacteria. Nat. Rev. Microbiol. 15, 621–629, (2017).

43. Pernthaler, J. Predation on prokaryotes in the water column and its ecological implications. Nat. Rev. Microbiol. 3, 537–546, (2005).

44. Matz, C. & Kjelleberg, S. Off the hook--how bacteria survive protozoan grazing. Trends Microbiol. 13, 302–307, (2005).

45. Silva, A. J., Leitch, G. J., Camilli, A. & Benitez, J. A. Contribution of hemagglutinin/protease and motility to the pathogenesis of El Tor biotype cholera. Infect. Immun. 74, 2072–2079, (2006).

46. Cinar, H. N. et al. Vibrio cholerae hemolysin is required for lethality, developmental delay, and intestinal vacuolation in *Caenorhabditis elegans*. PLoS One 5, e11558, (2010).

47. Yan, J., Nadell, C. D., Stone, H. A., Wingreen, N. S. & Bassler, B. L. Extracellularmatrix-mediated osmotic pressure drives *Vibrio cholerae* biofilm expansion and cheater exclusion. Nat. Commun. 8, 327, (2017).

48. Hatzios, S. K. et al. Chemoproteomic profiling of host and pathogen enzymes active in cholera. Nat. Chem. Biol. 12, 268–274, (2016).

49. Adiba, S., Nizak, C., van Baalen, M., Denamur, E. & Depaulis, F. From grazing resistance to pathogenesis: the coincidental evolution of virulence factors. PLoS One 5, e11882, (2010).

50. Yildiz, F. H. & Schoolnik, G. K. Role of *rpoS* in stress survival and virulence of *Vibrio cholerae*. J. Bacteriol. 180, 773–784, (1998).

51. De Souza Silva, O. & Blokesch, M. Genetic manipulation of *Vibrio cholerae* by combining natural transformation with FLP recombination. Plasmid 64, 186–195, (2010).

52. Marvig, R. L. & Blokesch, M. Natural transformation of *Vibrio cholerae* as a tool- optimizing the procedure. BMC Microbiol. 10, 155, (2010).

53. Blokesch, M. TransFLP - a method to genetically modify *V. cholerae* based on natural transformation and FLP-recombination. J. Vis. Exp. 68, e3761, (2012).

54. Borgeaud, S. & Blokesch, M. Overexpression of the *tcp* gene cluster using the T7 RNA polymerase/promoter system and natural transformation-mediated genetic engineering of *Vibrio cholerae*. PLoS One 8, e53952, (2013).

55. Gurung, I., Berry, J. L., Hall, A. M. J. & Pelicic, V. Cloning-independent markerless gene editing in *Streptococcus sanguinis*: novel insights in type IV pilus biology. Nucleic Acids Res. 45, e40, (2017).

56. Metzger, L. C. et al. Independent Regulation of Type VI Secretion in *Vibrio cholerae* by TfoX and TfoY. Cell Rep. 15, 951–958, (2016).

57. Miyazaki, K. Molecular engineering of a PheS counterselection marker for improved operating efficiency in *Escherichia coli*. Biotechniques 58, 86–88, (2015).

58. Bevis, B. J. & Glick, B. S. Rapidly maturing variants of the *Discosoma* red fluorescent protein (DsRed). Nat. Biotechnol. 20, 83–87, (2002).

59. Dunn, A. K., Millikan, D. S., Adin, D. M., Bose, J. L. & Stabb, E. V. New *rfp-* and pES213-derived tools for analyzing symbiotic *Vibrio fischeri* reveal patterns of infection and *lux* expression in situ. Appl. Environ. Microbiol. 72, 802–810, (2006).

60. Bao, Y., Lies, D. P., Fu, H. & Roberts, G. P. An improved Tn7-based system for the single-copy insertion of cloned genes into chromosomes of Gram-negative bacteria. Gene 109, 167–168, (1991).

61. Lo Scrudato, M. & Blokesch, M. The regulatory network of natural competence and transformation of *Vibrio cholerae*. PLoS Genet. 8, e1002778, (2012).

62. Meibom, K. L., Blokesch, M., Dolganov, N. A., Wu, C.-Y. & Schoolnik, G. K. Chitin induces natural competence in *Vibrio cholerae*. Science 310, 1824–1827, (2005).

63. Aubry, L., Klein, G., Martiel, J. L. & Satre, M. Kinetics of endosomal pH evolution in *Dictyostelium discoideum* amoebae. Study by fluorescence spectroscopy. J. Cell Sci. 105, 861–866, (1993).

64. Schneider, C. A., Rasband, W. S. & Eliceiri, K. W. NIH Image to ImageJ: 25 years of image analysis. Nat. Methods 9, 671–675, (2012).

65. Cardona, A. et al. TrakEM2 software for neural circuit reconstruction. PLoS One 7, e38011, (2012).

66. Schindelin, J. et al. Fiji: an open-source platform for biological-image analysis. Nat. Methods 9, 676–682, (2012).

